# Apical restriction of the planar cell polarity component VANGL in pancreatic ducts is required to maintain epithelial integrity

**DOI:** 10.1101/778332

**Authors:** Lydie Flasse, Siham Yennek, Cédric Cortijo, Irene Seijo Barandiaran, Marine R-C Kraus, Anne Grapin-Botton

## Abstract

Cell polarity is essential for the architecture and function of numerous epithelial tissues. Here we show how planar cell polarity (PCP), so far studied principally in flat epithelia, is deployed during the morphogenesis of a tubular organ. Using the mammalian pancreas as a model, we report that components of the core PCP pathway such as the transmembrane protein Van Gogh-like (VANGL), are progressively apically-restricted. VANGL expression becomes asymmetrically localized at the apical surface of ductal cells, revealing a planar polarization of the pancreatic duct. We further show that restricting VANGL to these discrete sites of expression is crucial for epithelial integrity. Expansion of expression on basolateral membranes of the progenitors leads to their death and extrusion from the epithelium, as previously observed for perturbations of apico-basal polarity. Using organoids and *in vivo* analyses, we show that cell elimination is induced by a decrease of Rock activity *via* Dishevelled.

## INTRODUCTION

Establishment of polarity is essential for the function of many cell types. In epithelia two axes of polarity have been characterized: 1) apico-basal polarity orients cells from the free surface or the lumen to the basal lamina and 2) planar polarity coordinates polarization of cell structures or behaviors in the plane of the tissue, orthogonal to the apico-basal axis. While visualizing the apico-basal axis is relatively easy, few cell types present overtly polarized structures suitable for the study of planar polarity. Much of what is known regarding planar cell polarity (PCP) has been gleaned from studies of flat monolayers in which planar polarity coordinates the orientation of cell appendices such as trichomes in fly wings or stereocilia bundles in the inner ear^1^. Here, we investigate its organization and function in more complex epithelia, such as tubes.

A key feature of planar polarity is the asymmetric localization of core molecules of the pathway along both the apical-basal polarity axis and the plane of the epithelium. An evolutionarily-conserved core group of six families of proteins is required to coordinate planar polarization between neighboring cells: the transmembrane proteins Frizzled (vertebrate: FZD; fly: FZ), CELSR (CELSR; FMI) and Van Gogh-like (VANGL; STBM/VANG) and the cytosolic-submembrane proteins Dishevelled (DVL; DSH), Inversin/Diversin (INV/ANKRD6; DGO), and Prickle (PK). The PCP signaling system employs intra- and inter-cellular feedback interactions between its core components to establish their characteristic asymmetric cellular distributions. In the most comprehensively-characterized system, the *Drosophila* wing blade, the FZ-DSH-DGO complex localizes to intercellular junctions facing the VANG-PK complex in the adjacent cell. FMI, an atypical cadherin, localizes to both sides of the apical junction and bridges the FZ and VANG complexes between neighboring cells. These intercellular interactions are counterbalanced intracellularly by reciprocal inhibitory interactions between the two complexes^1, 2^.

While traditionally depicted in flat epithelia that require directional motion or an oriented structure, it is less clear how these proteins are organized or function in complex epithelia forming tubular structures^3^. The developing pancreas consists of a network of tubes lined by polarized progenitors, the apical membranes of which line the duct lumen^4, 5^. This network is dynamic and remodeled throughout development while the pancreatic epithelium grows and progenitors commit to various specialized fates^6^. At embryonic day E10.5, multipotent progenitors are organized around discrete micro-lumens^7, 8^. By E12.5, these micro-lumens have fused to form a mesh of interconnected ducts. Distal “Tip” progenitors committed to an exocrine fate are restricted to the periphery while the core of this plexus contains bipotent “Trunk” progenitors that subsequently either remain in the epithelium to form mature duct cells or delaminate to give rise to endocrine cells. By birth, the ducts are remodeled into an arborized structure, transporting duct- and acinar-secreted bicarbonate and digestive enzymes respectively to the duodenum, while the endocrine cells coalesce into islets of Langerhans embedded near the ductal system.

As endocrine differentiation is decreased as a result of mutation of CELSR2/3, it suggests PCP components are essential regulators linking pancreas morphogenesis with cell fate^9^. However, how would these components be localized in a radially symmetric tubular epithelium and what would their function be in in tubular organs remains to be determined. VANGL, a component specific to PCP, plays a central role in the establishment of the core protein complex^10^. Mutations in the VANGLs underlie a broad spectrum of developmental disorders in humans^11^ such as neural tube defects^12, 13^ and renal^14, 15^, heart^16,17^ and lung diseases^18^. Moreover, recent studies have shown that VANGL2 is consistently upregulated and amplified in breast, ovarian and uterine carcinomas^10^ and a tumorigenic function has been established in rhabdomyosarcoma^19^.

In this study, we investigated how planar polarity is deployed in the development of a complex tubular organ, the pancreas, and how that spatial information is integrated to enable proper morphogenesis. We show that expression of the core PCP molecule VANGL2 is planar polarized and we highlight an enrichment of that protein at tricellular junctions. By perturbing the localization of VANGL2, we demonstrate that a stringent regulation of the level and localization of this protein at the membrane of progenitors is required to maintain epithelial integrity. In particular, we show that apical restriction of VANGL is essential for progenitor survival as an extension of its expression domain on the basolateral membrane leads to apoptosis *via* downregulation of the ROCK pathway. Moreover, systematic examination of the localization of other core PCP components reveals a cell-autonomous negative regulation of VANGL on Dishevelled, reinforcing the model of inhibitory intracellular interaction currently proposed in lower vertebrates.

## RESULTS

### Planar polarization of PCP components in the tubular epithelium of the pancreas

We have previously shown that the expression of VANGL1/2 is initiated in the pancreatic epithelium at around E11 when apico-basal polarity is first established^9^. Its cellular localization becomes progressively restricted to apical foci at cell junctions (Fig. 1a,b, S1d). In order to determine whether VANGL1/2 is asymmetrically expressed in the plane of the epithelium, we performed whole mount staining which allowed us to visualize pancreatic ducts in three dimensions (3D) (Fig. 1e-g). Co-staining for Mucin1 and β-Catenin, respectively marking apical and lateral membranes, revealed that ductal cells were anisotropic, exhibiting elongated apical surfaces along the longitudinal axis of the ducts (Fig. 1f,g). VANGL1/2 was distributed asymmetrically in the apical plane, the protein being enriched on the transverse membranes (perpendicular to the longitudinal axis of the duct) while it was almost absent on the longitudinal membranes (Fig. 1f,h and movie 1). While an asymmetric expression pattern characterizes the localization of PCP core components in flat monolayered epithelia, these findings reveal that this is conserved in a branched, tubular network. We also observed an enrichment of VANGL1/2 protein at tricellular junctions, “hot-spots” of epithelial tension^20^ (Fig. 1g).

**Figure 1.**
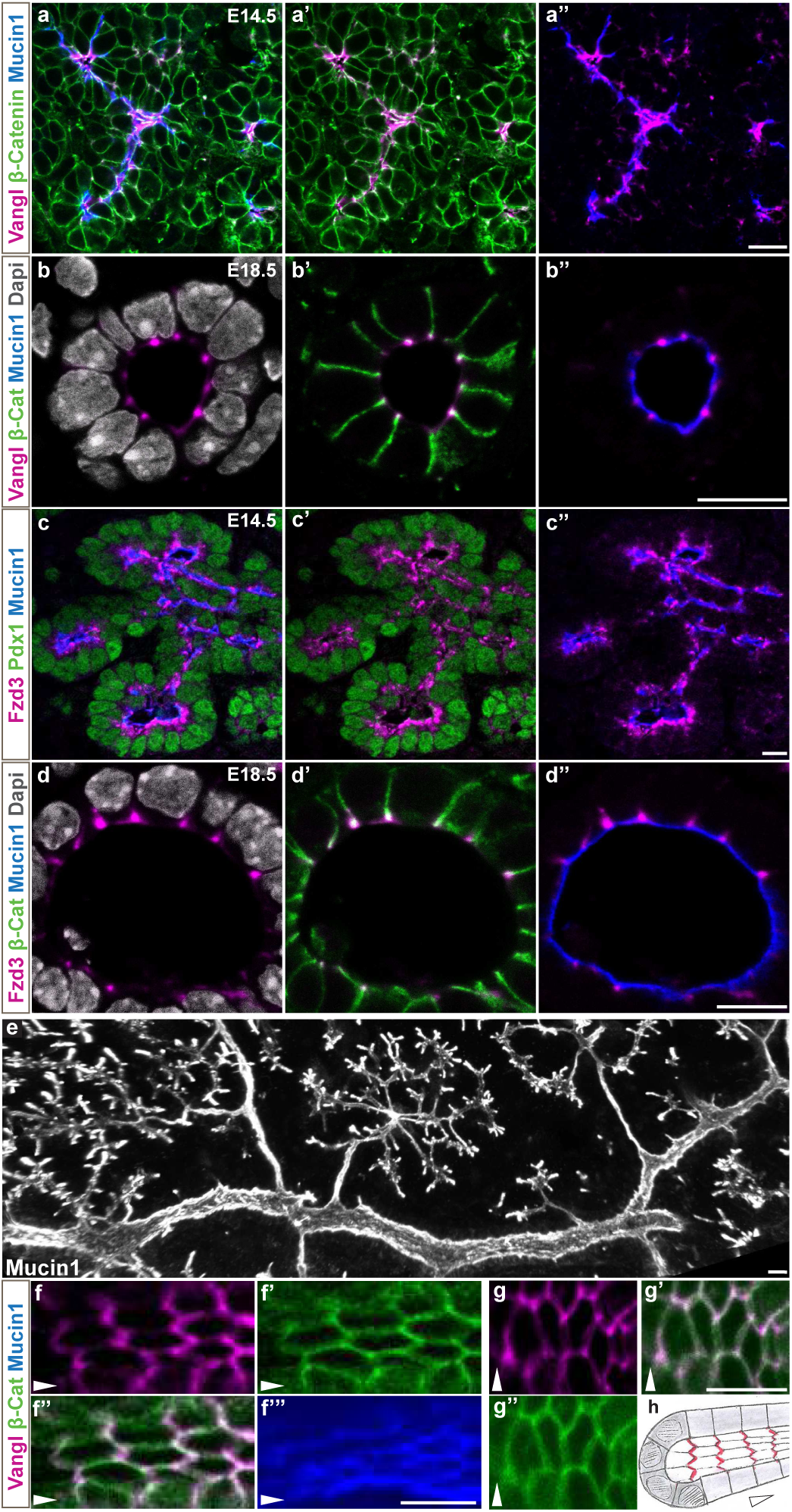
Planar polarization of PCP components in the tubular epithelium of the pancreas. Confocal images of embryonic pancreata at E14.5 (**a, c**) or E18.5 (**b, d-g**). β-Catenin labels all the membranes and Mucin1 the apical side. **a-d**, Representative images of VANGL (a-b) and FZD3 (c-d) expression, in a general view of the epithelium (a, c) or a close up of a transverse section of a duct (b, d), showing expression of those proteins at the apical cell junctions. **e**, 3D projection of a whole mount staining of Mucin1 depicting the architecture of the pancreatic duct tree. **f-g**, Confocal section of whole mount staining for VANGL showing representative apical views of the epithelium in a large duct, the arrowhead is pointing along the longitudinal axis of the duct. f illustrates the chevron-like localization of VANGL protein on the transverse membranes revealing the planar polarized organization of this protein. g illustrates the enrichment of VANGL protein at the tricellular junctions. **h**, scheme depicting VANGL (red) planar polarization in pancreatic ducts. Additional representation of VANGL expression pattern is shown in **movie 1**. Scale bar 10 µm.

We then investigated the expression of Frizzled proteins which is also planar polarized in flat epithelia. Although both FZD3 and FZD6 receptors can mediate PCP signaling^21^, we were only able to detect FZD3. Similarly to VANGL1/2 expression, at E10.5, small foci of FZD3 protein were detected around forming micro-lumens (Fig. S1a). At E12.5, FZD3 expression was enriched apically in the branching epithelium (Fig. S1b). Though largely restricted to the membrane, some cytoplasmic expression was visible (Fig. S1b). By E14.5, FZD3 was restricted to the apical-most regions of the lateral membranes, appearing as sub-apical striations of expression (Fig. S1c), which, by E18.5, condensed to foci at the apical cell junctions (Fig. 1d). Both FZD3 and VANGL1/2 were expressed in distal tip and proximal trunk cells of the epithelium, with VANGL1/2 expression becoming enriched in the trunk at later stages (Fig. 1a,c). Both were subsequently detected in islets of Langerhans at E18.5 (data not shown). Taken together, the membranous proteins of the PCP display progressive apical restriction in tubes as well as planar expression for at least VANGL.

### Ectopic expression of VANGL2 on baso-lateral membranes leads to cell death and pancreatic hypoplasia

Though the importance of asymmetric expression of PCP proteins in the plane is well established in both *Drosophila*^22^ and vertebrates^23^, little is known regarding their apical restriction. To test the significance of VANGL2 expression and localization, we generated transgenic mouse lines in which Cre-mediated recombination inactivates β-actin promoter-driven GFP expression and induces expression of a Cherry-VANGL2 fusion protein (Fig. 2d). This construct was previously reported to express a functional protein^24^. When expressed in pancreatic progenitors using *Pdx1*-Cre, the fusion protein was initially detected at the cell cortex in of most pancreatic epithelial cells at E10.5 and by E12.5 became membrane-enriched (Fig. S2a-b). Mesenchymal cells surrounding the epithelium expressed GFP, as expected from the non-recombined transgene (Fig. S2c). In contrast to endogenous VANGL proteins which are restricted to the apical junction in *wild-type* pancreata (Fig. 2a), expression of the Cherry-VANGL2 fusion protein showed no such delimitation and was detected throughout the membrane (Fig. 2b-c).

**Figure 2.**
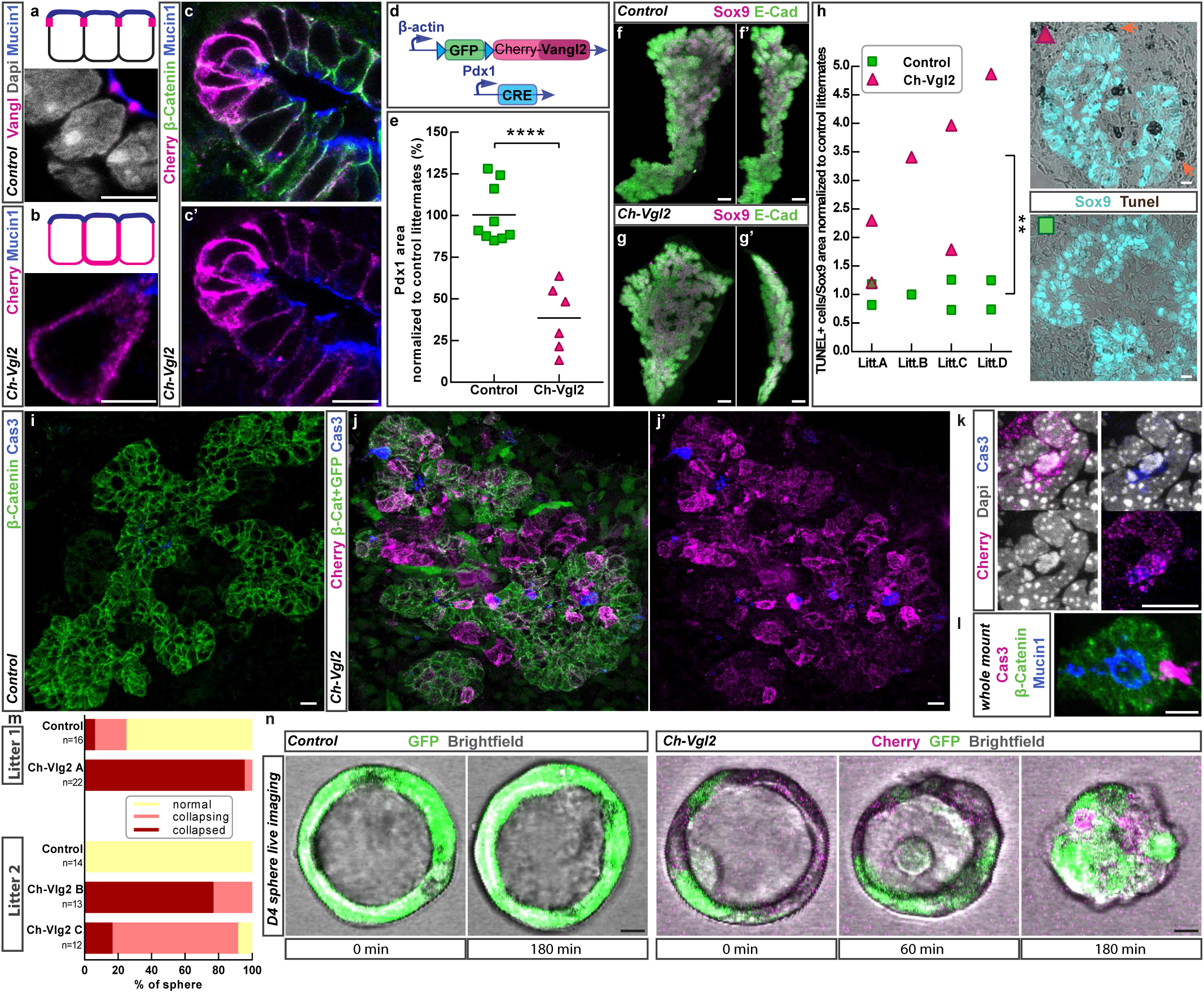
Ectopic expression of VANGL2 on baso-lateral membranes leads to disruption of epithelium integrity, cell death and pancreatic hypoplasia. **a-c**, confocal images showing localization of VANGL in a WT (a) or double-transgenic *Ch-Vlg2* (*Pdx1-Cre; Cherry-Vangl2*) (b) on an E16.5 pancreatic section. **d**, illustration of the construct used to express the fusion protein Cherry-VANGL2 in the pancreatic progenitors. **e**, PDX1 area quantified on E12.5 embryonic sections of control (green) and *Ch-Vlg2* (pink) pancreas showing a significant decrease of the epithelium size in the double-transgenic *Ch-Vgl2*. Each dot represents one pancreas, data normalized against control littermate. (****) p-value <0.0001 by t-test. **f-g**, whole mount immunostaining on E14.5 dorsal pancreas labelling progenitors (SOX9) and epithelial membrane (E-CAD). **h**, TUNEL assay labelling apoptotic cells in pancreatic epithelium (SOX9+) sections showing a significant increase of cell death in *Ch-Vlg2*. The ratios (number of TUNEL+ cells/epithelium area) are normalized against the control littermates and data are presented by litters. Each dot represents one pancreas. (**) p-value=0.004 by t-test. The orange arrow points at dead cells that left the epithelial layer. **i-k**, Caspase3 immuno staining on E12.5 pancreas sections showing dying cells in Cherry+ areas. The close-up in **k** shows one dying cell (CAS3+ and fragmented nuclei) expressing Cherry-VANGL2 protein. **l**, representative area of whole mount imunotaining illustrating delamination of a CAS3+ cell. **m-n**, pancreatospheres generated from control or *Ch-Vgl2* pancreata were imaged 4 days after seeding for either 3 hours (litter1) or 5 hours (litter2). **m**, Phenotypes were scored at the end of the movie: complete loss of lumen (collapsed), reduction of lumen size (collapsing), lumen diameter unchanged or increased (normal). A Chi-square test shows a significant difference between spheres generated from control or *Ch-Vgl2* with respect to the phenotype ‘reduced lumen size’ (collapsing + collapsed). Chi-square value = 52.6, p-value < 0.00001. **n**, Snapshots of representative spheres at the beginning and end of the movie. Additional examples of collapsing spheres are shown in **movie 2-7**. Scale bar 5 µm (a,b); 10 µm (c,h-l,n); 50 µm (f-g)

To determine whether this ectopic expression of VANGL protein throughout the cell membrane impacted upon pancreas development, we analyzed pancreatic epithelial size in double-transgenic *Pdx1-Cre; Cherry-Vangl2* (*Ch-Vgl2*) mice *versus* control (single-transgenic) littermates. At E10.5, when Cherry-VANGL2 protein was first detectable in double-transgenic animals, their pancreata were equivalent in size to controls (Fig. S2d) but by E12.5, we observed a ∼60% decrease in *Ch-Vgl2* pancreatic epithelial size (Fig. 2e, S2e). Pancreatic hypoplasia in *Ch-Vgl2* embryos was maintained at E14.5 (Fig. 2f,g) and was exacerbated at E16.5 (Fig. S2f). At this stage, we observed an ∼85% decrease in PDX1^+^ progenitors and endocrine (Glucagon^+^ or Insulin^+^) cells (Fig. S2f). Pancreatic hypoplasia was consistently observed in double-transgenic embryos generated from three independent lines of *Cherry-Vangl2* mice with varying transgene expression levels (Fig. S2e). Of note, for each line, the variability in pancreatic size between distinct *Ch-Vgl2* embryos likely reflects heterogeneity observed in the recombination rate between individuals.

To test whether hypoplasia of Cherry-VANGL2-expressing pancreata resulted from reduced proliferation, we quantified the proportions of phospho-HistoneH3-expressing cells in the epithelium. No significant difference was detected between the ratios of proliferating epithelial progenitors in E12.5 *Ch-Vgl2 versus* control siblings (Fig. S2g-h). In contrast, while TUNEL assays performed on E12.5 litters revealed only a few (0.07% in average) apoptotic cells in control pancreatic epithelia, we observed a 2.2-fold increase in TUNEL^+^ pancreatic epithelial cells in *Ch-Vgl2* littermates (Fig. 2h). Moreover, the extent of hypoplasia positively correlated with the proportion of apoptotic cells (Pearson correlation test: r=0.8; *p*<0.05) (Fig. S2k). In the *Ch-Vgl2* embryos, we also observed elevated TUNEL signal among the mesenchymal cells located just in the periphery of the epithelium (Fig. 2h, arrow and Fig. S2j). We presume that these cells, located at a distance of one or two cell diameters from the epithelium, are epithelial cells that have recombined and extruded from the distal-most epithelial layer. Staining for Caspase3 confirmed these observations, revealing an increased number of Caspase3^+^ cells in the epithelium expressing the Cherry-VANGL2 fusion protein (Fig. 2i,j), with some of those cells departing the outer-most epithelium (Fig. 2l). Furthermore, we detected dying (Caspase3^+^) cells retaining Cherry expression, suggesting that Cherry-VANGL2 expression cell-autonomously manifests in cell death (Fig. 2k). Taken together, these data show that ectopic expression of Cherry-VANGL2 protein in the pancreatic epithelium leads to apoptosis of the progenitors and, subsequently, to hypoplasia of the pancreas.

### Epithelial exit and destruction of epithelial integrity upon VANGL mislocalization to all cell membranes

To test the hypothesis that VANGL extension to all cell membranes may promotes epithelial exit, we conducted live imaging on pancreatospheres. These structures comprise a spherical monolayered pancreatic progenitor epithelium and are thus simpler than the branched pancreatic epithelium, and more amenable for observation of epithelial exit in live imaging^25, 26^. We generated pancreatospheres from *Ch-Vgl2* and control littermate pancreatic epithelium at E13.5, after PCP was established. We consistently observed increased cell delamination in spheres originating from *Ch-Vgl2* pancreata (movies 2, 3). These events lead to a disruption of the epithelium, resulting in inward collapse of pancreatospheres (Fig. 2m, n and movies 4, 5, 6). We have recently shown that pancreatic progenitors lining ducts secrete fluid into the lumen from E10.5^6^. We interpret that disruption of epithelium integrity leads to collapse because of either a relase of internal fluid pressure or disruption of supracellular apical cortical tension.

### VANGL misexpression leads to decreased actomyosin contractility

Actomyosin contraction, mediated by activation of Rho kinase, is well-recognised as an important downstream effector of the PCP^11, 27, 28^ and VANGL, in particular, has been involved in remodeling of actin microfilaments^29–33^. Moreover, actomyosin contractility is associated with apoptosis in different contexts^34^. For these reasons, we examined phosphorylation of myosin regulatory light chain (pMLC), a substrate of Rho kinase and effector/read-out of actomyosin-based contraction^35^ in pancreata from *Ch-Vgl2*. Immunostaining showed pMLC to be apically polarized in E12.5 control pancreata while the protein was undetectable in *Ch-Vgl2* pancreatic epithelia (Fig. S3a-b). At E14.5, pMLC was expressed at the apical cell junction in controls (Fig. 3a,c), mirroring the distribution of core PCP components, but pMLC was undetectable in Cherry-VANGL2-expressing epithelial domains (Fig. 3b). Western blotting on E12.5 (Fig. S3c) and E14.5 (Fig. 3d-e) dorsal pancreata confirmed these observations, revealing a 60% decrease in average of pMLC expression in E14.5 pancreata from *Ch-Vgl2* embryos compared with control siblings (Fig. 3e). Taken together, these results show a decrease of the ROCK pathway activity following ectopic VANGL expression.

**Figure 3.**
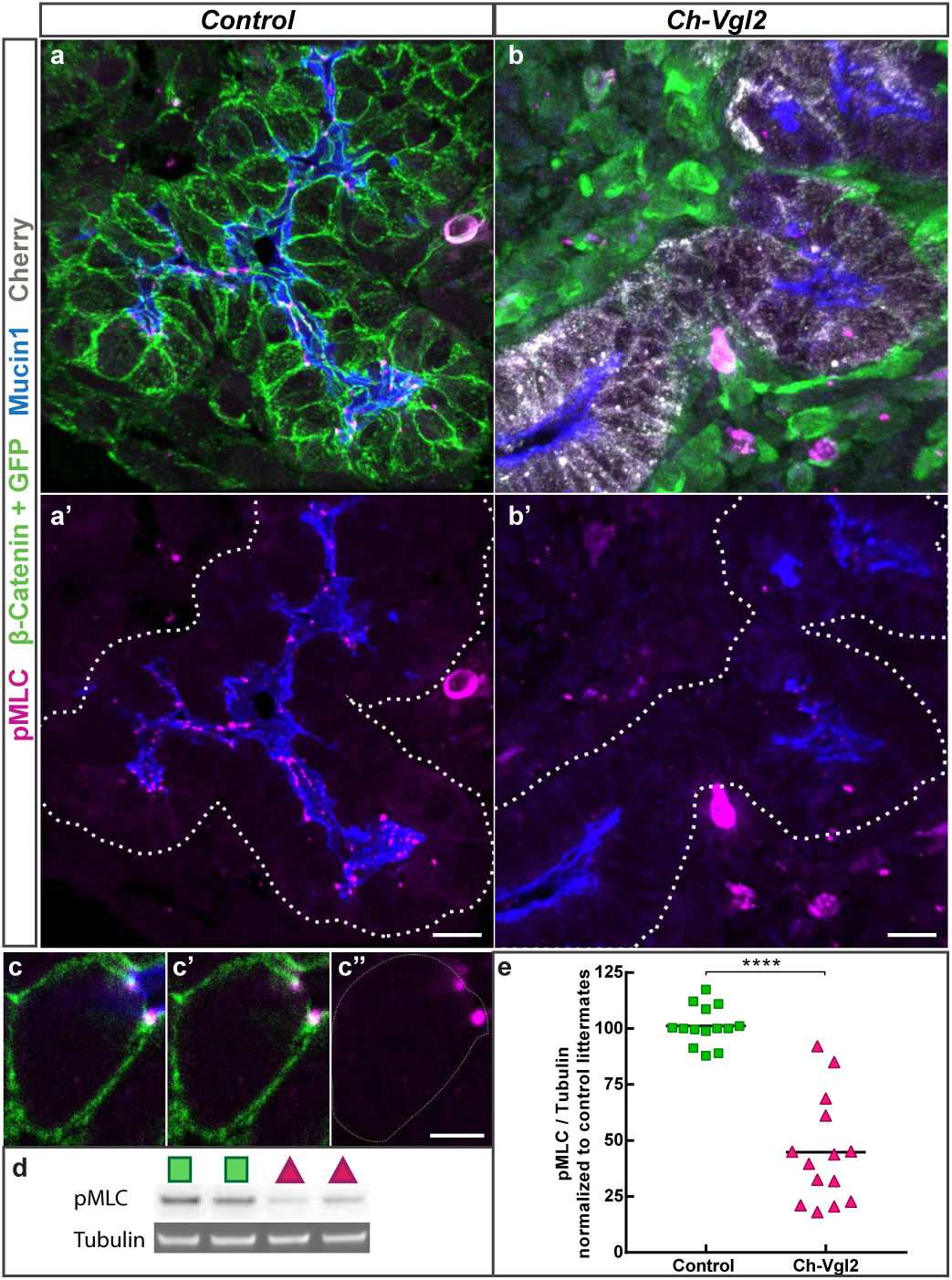
VANGL misexpression leads to a decrease of actomyosin contractility. **a-b**, 3D projection of confocal images showing expression of pMLC on E14.5 pancreas sections. Note the disappearance of pMLC apical foci in the *Ch-Vlg2* (*Pdx1-Cre; Cherry-Vangl2*) section (b). **c**, close up of a z section from the staining above highlighting the localization of pMLC protein at the apical cell junction in a WT sample. **d**, representative image of the pMLC western blots used for the quantification in e. **e**, pMLC/Tubulin ratios are normalized against the control littermates, each dot corresponds to one pancreas. Unpaired t-test reveals a significant decrease of pMLC protein level in the transgenic population compared to the control. (****) p-value <0.0001. Scale bar 5 µm (c); 10µm (a-b).

Jun N terminal Kinase (JNK) has been reported as a downstream effector of the PCP pathway although its function downstream of the core components and especially of VANGL protein remains unclear^36^. Western blots for phospho-c-jun did not reveal any significant dysregulation of the JNK pathway activity in E14.5 *Ch-Vgl2* compared with control pancreata (Fig. S3d-e). This suggests that the mechanism of VANGL misexpression-induced cell death is independent of the JNK pathway.

### RhoA pathway activation rescues cell death and pancreatic hypoplasia induced by VANGL misexpression

So far, our analysis showed an increase of cell death and a decrease of ROCK pathway activity following the misexpression of VANGL protein. In order to determine whether the decrease of ROCK pathway activity was responsible for apoptosis in *Ch-Vgl2* pancreata, we tested whether pharmacological activation of the ROCK pathway in *Ch-Vgl2* pancreatic epithelium is sufficient to rescue apoptosis and hypoplasia. To manipulate the ROCK pathway conveniently, we used an *ex vivo* culture system similar to our previously-published pancreatic organoid model^25^, but in which the whole E10.5 pancreatic epithelium was seeded in Matrigel enabling growth in 3D in the absence of mesenchyme. The growth of the developing pancreas was evaluated every day by measuring the circumference of the epithelial bud. The pancreatic buds expressing Cherry-VANGL2 protein grew slower and remained smaller than those from control littermates at the end of the culture period (five days), mimicking the phenotype observed *in vivo* (Fig. 4a,b). Inactivation of ROCK activity (*via* H1152, 5 µM) in *wild-type* epithelium manifested in an almost indistinguishable phenotype than the one observed for the *Ch-Vgl2* buds (Fig. 4c,d). Of note, we also observed a delay in branching of the pancreatic epithelium when the ROCK pathway was inhibited (Fig. 4c; asterisk). On the contrary, *wild-type* buds treated with a RhoA activator (CN03, 1 µg/ul) grew more rapidly and branched precociously compared with control vehicle-treated buds (Fig. 4e,f). Finally, activation of RhoA in *Ch-Vgl2* pancreatic epithelium rescued the hypoplasia (Fig. 4g,h, normalized data; for details, see *Materials & Methods*). In order to assess the level of activity of the ROCK pathway resulting from the various perturbations above, we performed Western blotting for pMLC on individual treated buds (Fig. 4i). While the amount of pMLC protein was decreased by ∼70% in untreated *Ch-Vgl2* buds (70 ± 4%) and *wild-type* buds treated with ROCK inhibitor (68 ± 7%), in *Ch-Vgl2* buds treated with RhoA activator, pMLC levels were comparable to controls, indicating that ROCK pathway activity was restored to endogenous levels (Fig. 4i). Lastly, we observed that *Ch-Vgl2* buds as well as those treated with ROCK inhibitor H1152 exhibited extensive apoptosis (Fig. 4j-l). In combination, these *ex vivo* experiments demonstrate that the cell death and pancreatic hypoplasia resulting from VANGL2 misexpression is a consequence of decreased ROCK pathway activity.

**Figure 4.**
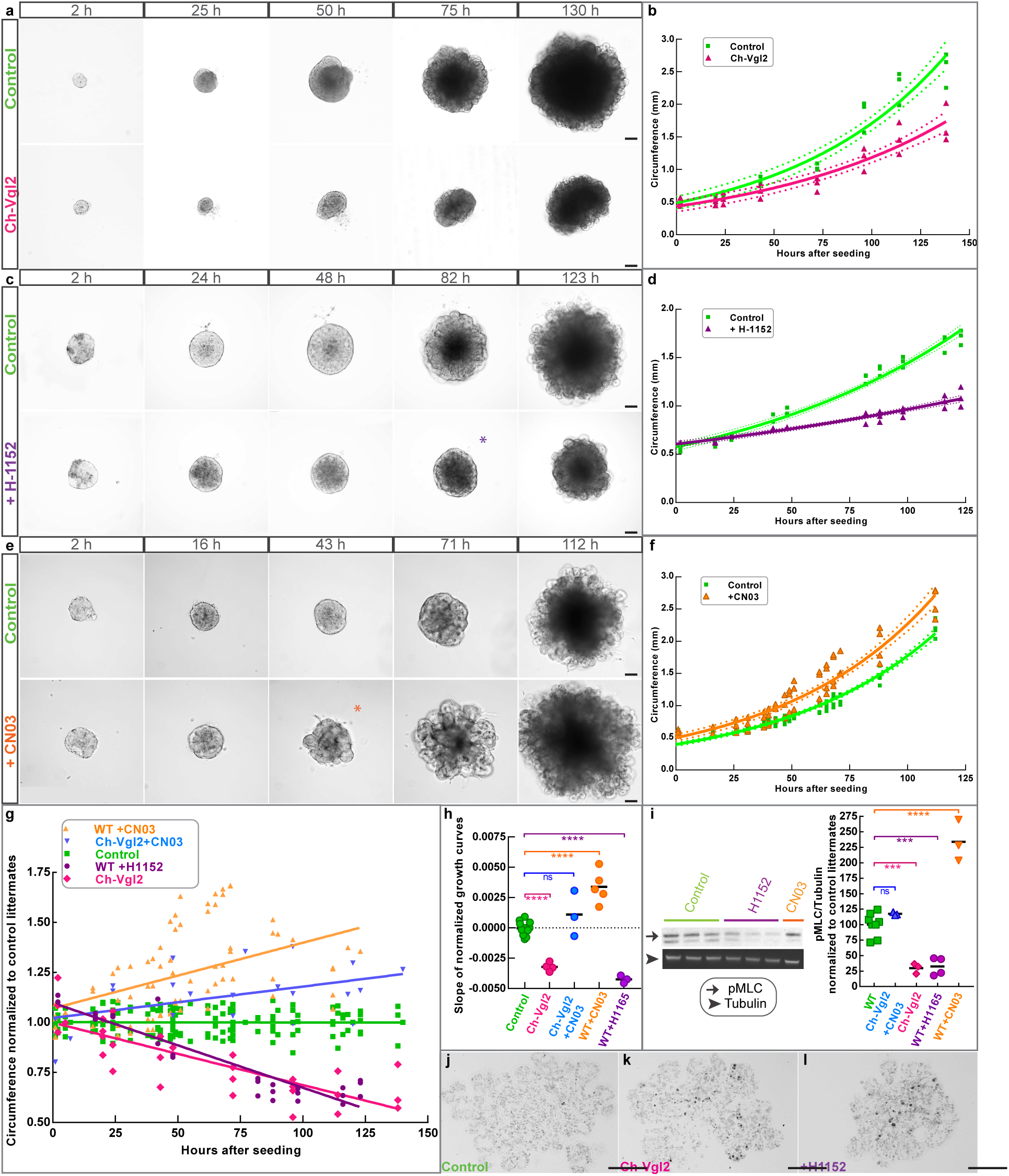
Activation of the RhoA pathway rescues cell death and hypoplasia induced by VANGL misexpression. **a-f**, Culture in 3D Matrigel of E10.5 pancreatic epithelium collected from WT or *Ch-Vgl2* (*Pdx1-Cre; Cherry-Vangl2*) embryos and submitted to treatment with ROCK inhibitor (H-1152) or RhoA activator (CNO3). **a, c, e**, pictures of the growing epithelium in each experimental condition at different time points of the culture (h=hours post seeding). **b, d, f**, graphs representing the quantification of the size of the epithelium, performed at each time point by measuring the circumference of the bud and fitting an exponential growth curve for each individual bud. For clarity, only an average growth curve is presented in each experimental condition. T-test computed on the doubling time of each curve demonstrates that double-transgenic (*Ch-Vgl2*, pink) and WT buds treated with ROCK inhibitor (purple) grow slower than their control littermates, p-value=0.01 and 0.006 respectively (n=3 in each condition). The buds treated with RhoA activator (orange) behave as controls (p-value= 0.8, n=3). T-test performed on the size of the bud at the end of the culture show that the *Ch-Vlg2* buds and the buds treated with ROCK inhibitor are smaller (p-value=0.02 and 0.001, respectively) while the bud treated with the activator became bigger (p-value=0.02). **g**, normalization of all the growth curves to the control littermates (see material and methods) showing that activation of the ROCK pathway in *Ch-Vlg2* can rescue the hypoplasia phenotype. **h**, statistical analysis computed on the slope of the normalized data presented in g. Each dot represents the growth of one bud; colors indicated the different experimental condition. (****) p-value <0.0001; ns=non-significant. **i**, representative image of a western blot against pMLC and quantification of the band intensities in the different experimental conditions showing that the rescued buds display similar level of Rock pathway activity relative to the control. One dot represents the pMLC/Tubulin ratio for an individual bud. Data were normalized to the control littermates. (****) p-value <0.0001; (***) p-value ≤0.0005; ns=non-significant. **j-l**, TUNEL assay on bud sections shows an increase of apoptosis in the *Ch-Vgl2* buds (k) as well as in the bud treated with ROCK inhibitor (l) compared to the control sibling (j). Scale bar 100 µm.

### Ectopic VANGL expression interferes with Dishevelled but not with Prickle or Frizzled localization

VANGL physically interacts with the majority of the other PCP components^23^ and, depending on the context, manipulation of VANGL expression can affect the level and/or the localization of these proteins. Hence, we systematically examined the expression pattern of VANGL2 partners possibly involved in the downregulation of actomyosin contractility.

Frizzled and VANGL interact across the membranes of adjacent cells, bridged by the atypical cadherin CELSR, enabling the propagation of polarity from cell to cell. In the *Ch-Vgl2* pancreas, Frizzled3 expression remained at the apical junction and the expression level of the protein was unaffected (Fig. 5a-d). Thus, VANGL basolateral expansion fails to induce a similar expansion of Frizzled3. This suggests that intercellular communication is not drastically dysregulated following ectopic VANGL expression, as Frizzled receptor in the neighboring cells acts as a limiting factor.

**Figure 5.**
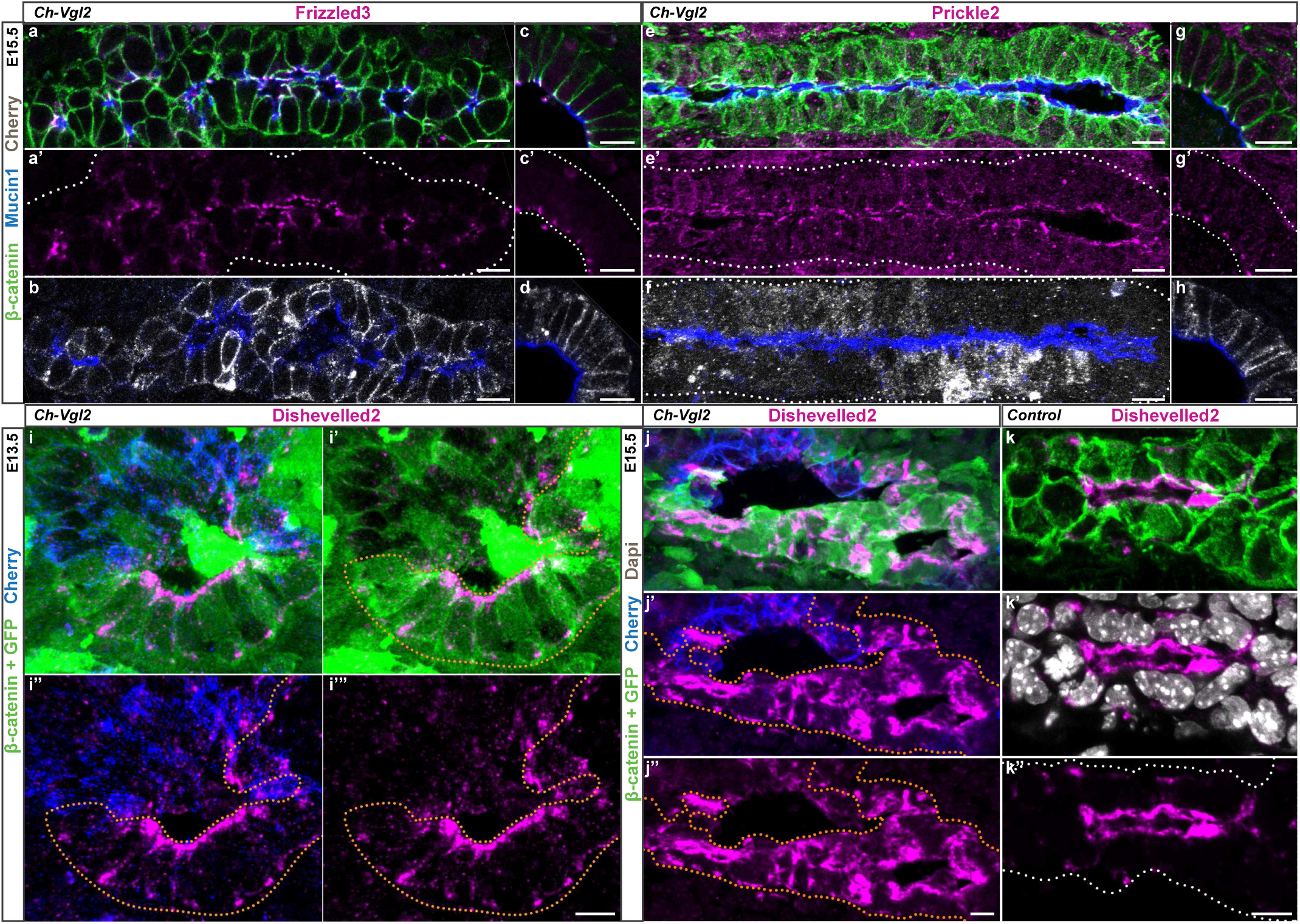
Ectopic VANGL expression interferes with Dishevelled but not with Prickle nor Frizzled localization. Confocal images of E15.5 (a-h, j-k) or E13.5 (i) pancreatic sections, showing the expression pattern of PCP components in Cherry-VANGL2+ area. (a-d, g-h) Z-sections and (e-f, i-k) 3D projections are displayed. Longitudinal (a-b; e-f) and transversal (c-d; g-h; i-k) sections of pancreatic ducts. **a-d**, Frizzled3 (FZD3) is expressed at the apical cell junctions in the area expressing the fusion protein Cherry-VANGL2 showing that VANGL basolateral expansion fails to induce a similar expansion of FZD3. Section b and d are adjacent to a and c respectively. **e-h**, Prickle2 (PK2) is expressed at the apical cell junction in the Cherry+ area. No extension of PK2 expression domain on the basolateral membrane is observed. Sections f and h are adjacent to e and g respectively. **i-k**, Dishevelled2 expression (magenta) is decreased in the Cherry+ cells (blue) compared to the adjacent cells that are not expressing the Cherry-VANGL2 protein (GFP+ cell; green and highlighted by orange dots) or the WT cells (k). Scale bar 10 µm.

Intracellularly, Prickle directly binds to VANGL and can bind to Dishevelled to antagonize its Frizzled-mediated membrane recruitment^37–40^. Like VANGL1/2 and Frizzled3, Prickle2 is expressed at the apical junctions in the pancreatic epithelium (Fig. S4a-b). In *Ch-Vgl2*, we did not detect any changes in the pattern of expression of Prickle2 at E13.5 or E14.5 (Fig. 5e-h and data not shown).

Dishevelled can be physically bound by VANG in *Drosophila*, and in vertebrates, these interactions are believed to antagonize the formation of the FZD-DVL complex by affecting DSH/DVL levels and/or stability^28, 37, 41, 42^. In the E12.5 pancreatic epithelium, DVL2 protein is enriched apically in the duct and becomes cytoplasmic in the newly-formed endocrine clusters (Fig. S4c). This expression pattern is maintained at E15.5 (Fig. S4d-g). High resolution expression analysis at this stage showed that contrary to the other core components, DVL2 protein is not tightly restricted to the apical junction (Fig. S4g) but instead extends sub-apically, forming a ring just below the apical membrane (Fig. S4e). In *Ch-Vgl2* pancreata at E13.5 and E15.5, we observed a decrease in DVL2 expression in the cells expressing Cherry-VANGL2 protein (Fig. S4h; Fig. 5i-k). These results support the notion that ectopic expression of VANGL2 perturbs the level and/or stability of DVL2 in a cell-autonomous manner, which in turn leads to a decreased activity of the downstream effectors involved in actomyosin contractility.

Although VANGL is mainly known to regulate PCP, previous reports have shown that VANGL2 may be involved in maintaining apico-basal polarity^10, 31, 43–45^. Therefore, we investigated whether the extension of the VANGL2 expression domain to the basolateral membrane affects the localization of apico-basal markers. The cellular localization of the apical membrane glycoprotein Mucin1 (Fig. 5a-h), the apical junction marker PAR3, the tight junction protein ZO-1 and the basolateral marker Scribble are all unchanged in Cherry-VANGL2-expressing cells (Fig. S5a-e), indicating that the integrity of the apico-basal axis is maintained in *Ch-Vgl2* pancreata.

### Apical overexpression of VANGL protein due to inversin mutation does not perturb actomyosin contractility or induce cell death

Since the function of VANGL relies on its cellular localization, we sought to distinguish whether cell death and epithelial exit in *Ch-Vgl2* pancreata are due to the ectopic expression of VANGL on the basolateral membrane or overexpression of the protein in general. Diego and its distant vertebrate homologue Inversin, are known to bind directly with Van Gogh/VANGL protein^46, 47^ and genetically interfere with VANG function^47, 48^. Analysis of *the Inversin* mutant *(Invs^Inv/Inv^*)^47, 49, 50^ pancreata revealed increased levels of VANGL protein at the apical side of pancreatic ducts (Fig. 6a-d). However, conversely to our observations in *Ch-Vgl2*, we showed that the expression domain of VANGL did not extend to the baso-lateral membrane but rather to the apical membrane in the *Invs^Inv/Inv^* pancreas (Fig. 6b,d).

**Figure 6.**
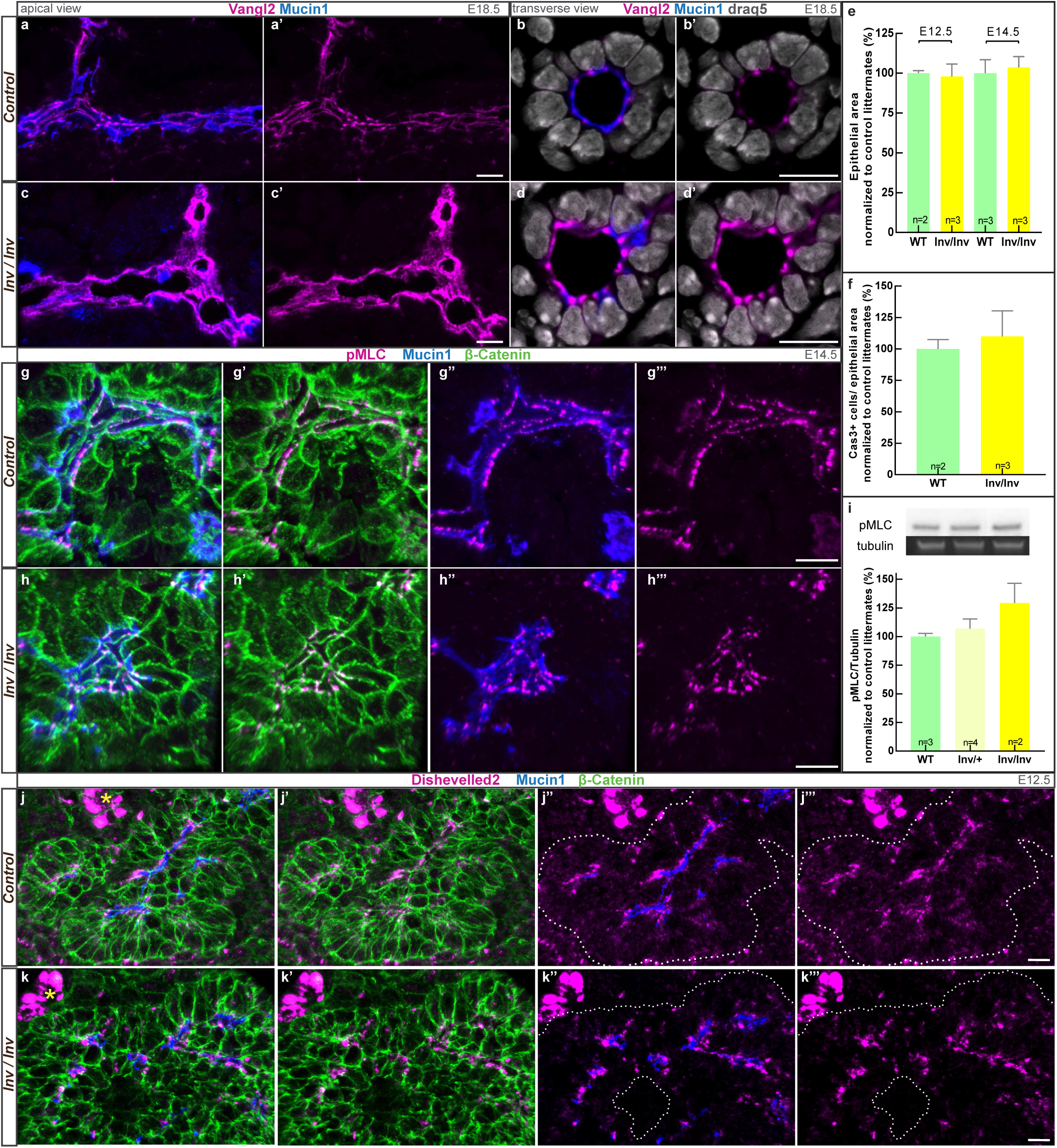
Apical overexpression of VANGL protein due to Inversin mutation does not lead to the perturbation of actomyosin contractility nor cell death. **a-d**, longitudinal (a, c) and transversal (b, d) sections of E18.5 pancreatic duct immunolabelled for VANGL (magenta) show an increase of the protein at the apical membrane in the inversin mutant (c, d) compared to the WT littermate (a, b). **e**, quantification of the epithelium area in the inversin mutant and the control littermates at E12.5 and E14.5 shows no difference in the size of the epithelium. p-value>0.5 by t-test. **f**, quantification of the number of Caspase3 positive (CAS3+) cells in E12.5 inversin mutant shows no significant difference in apoptotic cells compared to the control. The CAS3+ cells/ epithelium area ratio was normalized against the control littermates. p-value>0.5 by t-test. **g-h**, staining for pMLC in E14.5 WT and Inversin pancreatic sections shows no differences in the localization of the protein. **i**, representative image of a western blot against pMLC. Quantification of the pMLC/Tubulin ratio shows no difference between inversin homozygote, heterozygote mutant and their control littermates. p-value>0.1, by t-test. **j-k**, Z-projection of confocal images showing Dishevelled2 (DVL2) expression on E12.5 pancreatic sections. No differences of localization or level of DVL2 were observed between the control (j) and the inversin mutant (k). *, highlighted endocrine islet. Scale bar 10 µm.

This model of apical overexpression of VANGL without lateral expansion does not display any defect in the size of the pancreas at E12.5 and E14.5, nor increased apoptosis (Fig. 6e,f). Moreover, the level and the localization of pMLC were unchanged in *Invs^Inv/Inv^* mutant pancreata relative to controls (Fig. 6g-i), suggesting that the activity of the ROCK pathway is unaffected. Finally, DVL2 protein expression levels and localization were also unaffected in the *Invs^Inv/Inv^* pancreatic epithelium (Fig. 6j,k).

Overall, these findings show that while both *Invs^Inv/Inv^* and *Ch-Vgl2* exhibit overexpression of VANGL at the membrane, only ectopic expression of VANGL specifically on the basolateral membrane leads to cell death and epithelial exit. This strongly suggests that an apical restriction of VANGL protein activity is essential to maintain epithelial integrity.

### Misexpression of VANGL2 protein in the neural tube also induces apoptosis

In order to determine whether we could extend our findings to other epithelia besides the pancreas, we induced the expression of Cherry-VANGL2 protein in the neural tube using the *Rosa26*^CreER^ driver. VANGL protein is polarized apically in this tissue and plays an essential role in neural tube closure and elongation^51^. Induction of the transgene was performed at E11 by injection of 4OH-tamoxifen into the pregnant female and embryos were harvested at either 6, 14, 24 or 48 hours post-injection. Staining for Caspase3 revealed a significant 3.3-fold increase in the proportion of apoptotic cells within the neural epithelium following the induction of the transgene, peaking at 14 hours post-injection (Fig. 7a-c). Therefore, we can conclude that a VANGL misexpression induces apoptosis not only in pancreas but also in other epithelia such as the neural tube.

**Figure 7.**
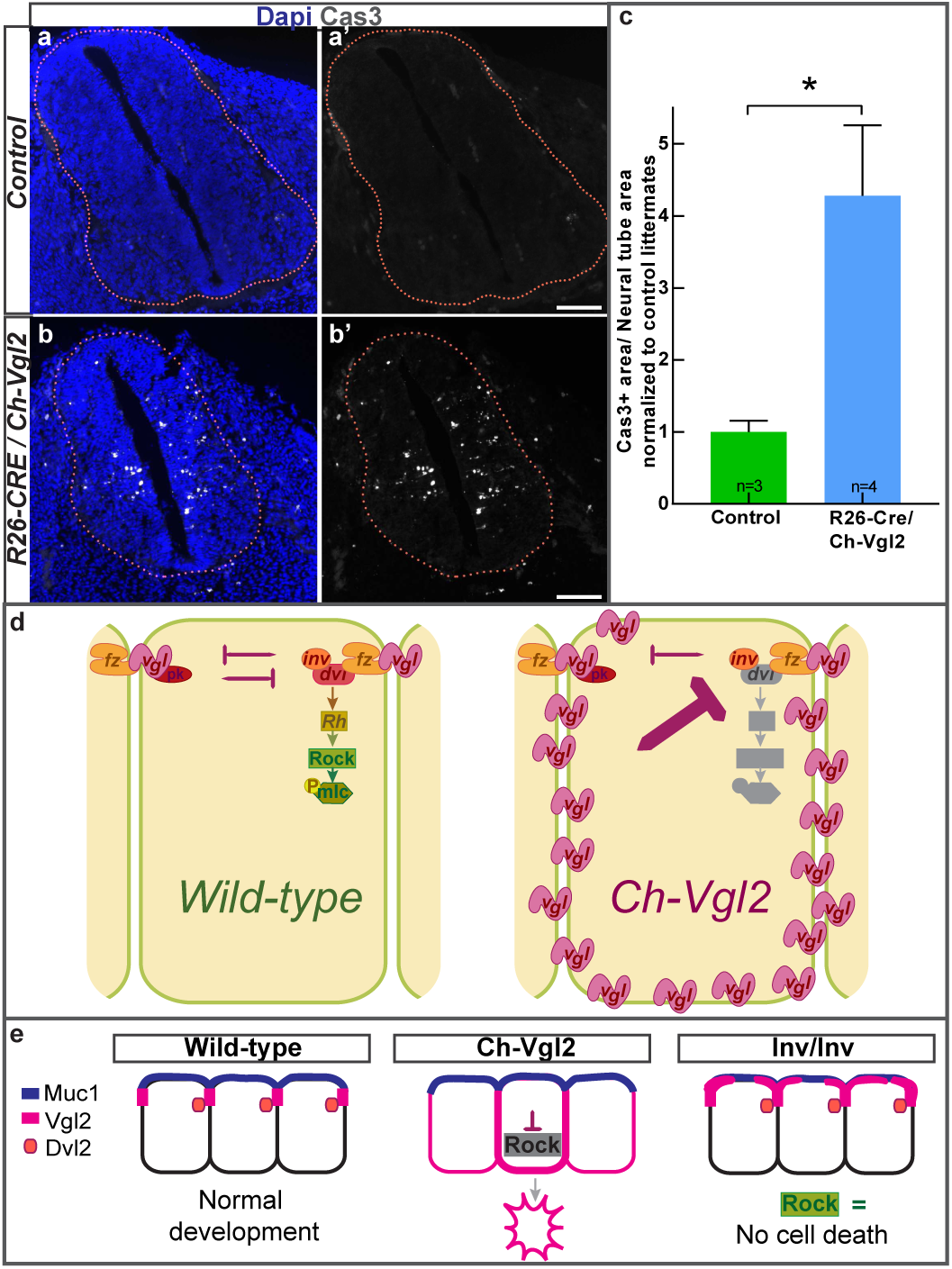
Misexpression of VANGL2 protein in the neural tube also induces apoptosis. **a,b**, representative images of Caspase3 immunostaining on whole embryo sections showing an increase of apoptotic cell in the neural tube of *Rosa26*^CreER^; *Cherry-Vangl2* embryos collected 14h after induction of transgene expression relative to control (*Rosa26*^CreER^). **c**, Quantification of the ratio CAS3^+^ area/neural tube area shows a 3.3 fold increase of apoptosis following expression of Cherry-VANGL2 protein. (*) p-value <0.05 by t-test. Scale bar 100 µm. **d**, Working model: in wild-type cells (left) VANGL expression is restricted to the apical junctions and the Rock pathway is activated downstream of the FZD/DVL complex. In the *Ch-Vgl2* double-transgenic (right) VANGL expression is extended to the latero/basal membranes and Rock pathway activity is decreased. Since FZD protein level is not increased at the apical cell junctions of the transgenic cells, the intercellular activation of the distal FZD/DVL/INV complex by VANGL is not increased. Rather, cell autonomous misexpression of VANGL leads to a decrease of DVL protein and consequently of the downstream ROCK activity. **e**, Results summary.

## DISCUSSION

In this study we elucidate how planar polarity is deployed in a complex tubular organ and we demonstrate that proper localization of the PCP core component VANGL is required for survival of pancreatic progenitors and, hence, for normal pancreatic morphogenesis. Using transgenic lines driving the expression of Cherry-VANGL2 protein all around the membrane of pancreatic progenitors, we show that ectopic expression of this protein in the basolateral domain leads to apoptosis *via* downregulation of the ROCK pathway. Live imaging of pancreatospheres revealed this phenomenon to be associated with a disruption of epithelial integrity. Moreover, systematic expression analyses of core PCP components allowed us to decipher a mechanism of intracellular inhibitory interactions of possible relevance to other organs. We propose a model whereby extension of the VANGL membrane expression domain increases the cell-autonomous repression exerted on Dishevelled, hence preventing its localization at the apical membrane (Fig. 7d). As VANGL can bind Dishevelled in the fly^37^ as well as in vertebrates^41^, VANGL may directly destabilize DVL and inhibit its recruitment to the junctions by FZD.

This hypothesis is strongly supported by the work of the Tomlin lab in *Drosophila* showing the ability of VANG to cell-autonomously block recruitment of DSH at the membrane^52^, resembling the previously-reported repression of DSH by PK^39, 40, 52^. In our system, an expansion of Prickle does not result from misexpression of VANGL protein and is therefore unlikely to contribute to the repression exerted on DVL. Recruitment of Dishevelled at the membrane is essential for its function in PCP^53, 54^. More specifically, its membrane localization in *Xenopus* gastrulating cells is required for activation of Rho and Rac^55^, the small GTPases active downstream of the Frizzled-Dishevelled module. Our results corroborate these data in mammals as we show a decrease of ROCK activity in Dishevelled^low^ (Cherry^+^) cells. Precisely how VANGL perturbs DVL level/stability remains unclear but it may operate *via* ubiquitin-mediated degradation, a mechanism recently proposed to modulate PCP protein levels^56, 57^. In particular VANGL1/2 promotes NRDP1-mediated Dishevelled ubiquitination in cell lines^58^. Interestingly, in the *Inversin* mutant which exhibits an upregulation of VANGL protein exclusively at the apical side, such a downregulation of Dishevelled protein is not observed. Neither increased cell death, epithelium integrity defects nor ROCK activity dysregulation were detected. Therefore, it appears that the ectopic localization of VANGL on the basolateral membrane is the key determinant in the mechanism we propose and that restriction of VANGL protein activity to the apical membrane is essential for cell survival.

Interactions between apico-basal polarity and planar polarity have been reported^11, 59^ but, so far, only apico-basal polarity defects have been associated with apoptosis. In the fly it is well established that disruption of apico-basal components such as crumbs, stardust or bazooka lead to apoptosis^60–62^. Our work hereby reveals that disruption of a PCP core component expression domain can also manifest in cell death. The downstream pathways involved differ as in fly, activation of the JNK pathway triggers apoptosis^63^ while we show that a decrease of ROCK pathway activity induces apoptosis following ectopic expression of VANGL. ROCK signaling can serve as either a pro-apoptotic or pro-survival cue in a cell type and context-dependent manner^64^. In particular, in airway epithelial cells, ROCK inhibition induces apoptosis *via* activation of the caspase cascade^65^. Our studies suggest that this is similarly operating in the pancreatic epithelium. We also show that Cherry-VANGL2-positive cells exit the epithelium, suggesting that ectopic VANGL expression promotes delamination in a cell-autonomous manner. The causal relationships between actomyosin dynamics, delamination and cell death have been debated in recent investigations^34^. Hence, one hypothesis is that mosaic transgene expression results in varying levels of actomysosin activity between cells, leading cells with low actomyosin activity to be excluded by neighboring cells exerting higher tension before or after cell death. Such mosaic patterns are reminiscent of cell competition processes observed in apico-basal polarity-deficient clones. In both fly and MDCK cells, cells lacking basolateral determinants such as Scribble, Mahjong and Lethal giant larvae undergo apoptosis if *wild-type* cells surround them^66^. Interestingly, Scribble-deficient clones are also eliminated by extrusion and, ultimately, apoptosis^67, 68^. These competition mechanisms which eliminate abnormal cells, are essential to prevent tumor development and such a mechanism may be at play in the case of abnormal VANGL-polarized cells. Vang Gogh-like proteins are upregulated in many types of aggressive cancers and this upregulation is associated with poor patient prognosis in breast cancer^69–71^.

In addition to revealing the crucial nature of PCP components in maintaining epithelial integrity in two tissues, using whole mount staining enabled us to visualize planar polarity in three dimensions in a complex ductular network. VANGL and other PCP proteins have been shown to control the development of tubular organs such as the branching of the lung^72^ or the diameter of the duct in the kidney^73, 3^ where planar-oriented proteins and cell behaviors have been inferred from 2D sections^74, 75^. Our study showing PCP organization in 3D reveals that a chevron organization of VANGL orients the cells along the longitudinal axis of the duct (Fig. 1h). This organization, the striking elongation of cells along the duct length and VANGL enrichment at tricellular junctions, known “hot-spots” of epithelial tension, are all suggestive of a link between PCP component localization and anisotropic stresses in tubes^20, 76^. To our knowledge, such enrichment of a PCP core component at tricellular junctions has never been reported.

As these specialized junctions are involved in cell translocation between epithelial layers or cell intercalation^20^, an increase of VANGL at the tricellular junctions may affect the delamination of endocrine progenitors, thereby linking morphogenesis/architecture to the endocrine differentiation program. Further work investigating the integrity of the ductal network as well as cellular differentiation in the absence of VANGL protein will be essential to answer these questions.

## Acknowledgements

We would like to thank Manuel Figueiredo-Larsen for his contribution to the experiments with whole-pancreas organoids. We thank Danelle Devenport for the *Cherry-Vangl2* construct, Nancy Thompson and the EPFL transgenesis facility for generating the transgenic lines. Thanks to J. Bulkescher and G. Karemore for their advice on imaging and statistics respectively and to Josh Brickman and Phil Seymour for their comments on the manuscript. Many thanks to Toshihisa Ohtsuka and Jeremy Nathans for the Prickle2 and Frizzled3 antibodies respectivly. The Novo Nordisk Foundation Center for Stem Cell Biology is supported by a Novo Nordisk Foundation grant number NNF17CC0027852

## Author contributions

L. F. contributed to project design as well as most experimental data and analyses; S. Y. contributed to all experiments with the ex vivo systems; I. S. B. contributed to the quantification of Caspase3^+^ cells in the inversin mutant and in the neural tube and participated to the analysis of the pancreatosphere movies; C.C. performed characterization of hypoplasia in *Cherry-Vangl2* line at E16.5; M. K. quantified pancreatic size of Inversin mutant at E14.5; C.C., M. K. and L.F established VANGL up-regulation in *Inv* mice. A. G-B. supervised the project and helped design the project. All authors contributed to drafting or revising the manuscript.

## METHODS

### Mice

Mice (Mus musculus) of mixed background were housed at the University of Copenhagen. All experiments were performed according to ethical guidelines approved by the Danish Animal Experiments Inspectorate (Dyreforsøgstilsynet). The following genetically modified mouse lines were used: *Gt(ROSA)26Sortm1(cre/ERT2)Tyj/J (Rosa26CreER)*^77^; *Invs*^Inv^ ^50^; *Tg(Ipf1-cre)1Tuv (Pdx1-CRE)*^78^. For the *Cherry-Vangl2* lines *(Tg (pCAGGS-LoxP-Cherry-Vgl2)AGB)* : the amino-terminally tagged *Cherry– Vangl2* fusion protein construct^24^ was inserted into a *CMV-CAG-loxP-eGFP-Stop-loxP-IRES-bGal* expression vector^79^. Three lines (7979; 9140 and 9139) expressing different level of the transgene were generated by random insertion, the two lines expressing the highest level of the Cherry-VANGL2 protein (7979 and 9140) were used to collect the data. *Pdx1-Cre; Cherry-Vangl2* double-transgenic embryos were obtained by crossing heterozygous *Pdx1-Cre* with heterozygous *Cherry-Vangl2* transgenic mice. After checking that the single transgenic did not present any phenotype, we used WT and single transgenic (*Pdx1-Cre* Tg/+ or *Cherry-Vangl2* Tg/+) indiscriminately as controls. Conditional induction of Cherry-VANGL2 protein was performed by intraperitoneal injection of 4-OH tamoxifen (4-OHTm, Sigma, H6278) at a concentration of 66,6 μg/g in *Rosa26CreER* females mated with *Cherry-Vangl2* males. The 4-OHTm solution was prepared following instructions described in Chevalier et al^.80^

### Whole-mount immunohistochemistry

E18.5 dorsal and ventral pancreata embedded in the first loop of duodenum were fixed in 4% paraformaldehyde (PFA) for 2h at room temperature (RT) while E14.5 dorsal pancreata were fixed 30 min or 1h at RT. After washing in PBS and stepwise dehydration in methanol (MeOH), fixed tissue was stored in 100% MeOH at −20°C. To quench autofluorescence, samples were incubated with freshly prepared MeOH:DMSO:H_2_O_2_ (2:1:3, 15% H_2_O_2_) for 12–24 hours at RT. Samples were washed twice in 100% MeOH for 30 minutes at RT. To improve permeability of the tissue, 3 cycles of incubation at –80°C were used (1h at −80°C followed by 1h at RT). Samples were rehydrated stepwise in TBST (50 mM Tris-HCl pH 7.4, 150 mM NaCl, 0.1% TritonX-100) 33%, 66%, and 100% for 15 min at each step at RT. For VANGL staining, an antigen retrieval step was performed: tissue was immerged in 10mM trisodium citrate buffer pH6.0 and brought, under low pressure setting, to 110°C for 15 min using the TintoRetriever system from BioSB. After blocking 24h at 4°C in TNB blocking solution (Perking Elmer TSA kit, NEL704A001KT) for VANGL staining or in CAS-Block (Thermo Fisher, #8120) for the other stainings, the specimen were incubated with primary antibodies for 48 hours at 4°C. Samples were washed extensively in TBST (at least 5 incubations of 15 min) and incubated with secondary antibodies (for VANGL staining, anti-rabbit biotinylated was used) for 48 hours at 4°C. After extensive washes in TBST, a 48h incubation with Streptavindin-HRP (Perking Elmer TSA kit) was performed to amplify VANGL signal. Other stainings performed in whole-mount did not require any amplification, the samples were therefore dehydrated stepwise in MeOH at this step. Samples stained with VANGL antibody were washed again in TBST before incubation with Cy3 for 1h at RT (Perking Elmer TSA kit). Then, after washing in TBST, the specimen were dehydrated in MeOH and stored at −20°C. All samples were cleared in a solution of 1:2 Benzyle Alcohol and Benzyle Benzoate (BABB) for 12–24 hours prior to imaging. Cleared specimens were subsequently mounted in glass concavity slides in BABB. Imaging was performed using a Leica SP8 confocal microscope with a 20x/0.75 IMM CORR objective and Hybrid detectors at 1024 × 1024 resolution in an 8-bit format. Three-dimensional reconstructions and movies were performed with the Imaris software (Bitplane).

### Immunohistochemistry on sections

The whole pancreas and the surrounding tissue (duodenum, stomach and spleen) were isolated in one piece and fixed in 4% PFA for 30min (E10.5 and E12.5), 1h (E14.5) or 2h (E18.5) at RT. E13.5 and E15.5 samples prepared for DVL2 staining were fixed 30 min in PFA diluted to 4% in water rather than the usual PBS. After several washes in PBS, samples were equilibrated in 15% sucrose/PBS 0.12 M solution overnight at 4°C prior to embedding in gelatin (7.5% gelatin diluted in 15% sucrose/PBS 0.12 M solution). Gelatin blocks were subsequently frozen at −65°C in 2-methylbutane (isopentane, Acros Organics) and kept at −80°C until sectioning. Frozen sections (8 to 10 µm) were then dried at room temperature and washed in PBS Triton 0.1%. The slides were incubated 10min in PBS Triton 0.3% to permeabilize the tissue. If necessary, an antigen retrieval step was performed at this point (see below). If secondary antibodies directly conjugated to HorseRadish Peroxidase (HRP) were used, a 7 min incubation in 3% H_2_O_2_ was performed at this point. The tissues were rinsed and then blocked 1h at RT in the appropriate blocking reagent (Table 1, supp. Methods). Primary antibodies were incubated overnight at 4°C (Table 2, supp. Methods). Following 3 washes in PBS Triton 0.1%, the samples were incubated with secondary antibodies for 1h at RT then rinsed again. For the antibodies requiring amplification (Table1, supp. Methods): if the secondary antibody was directly conjugated to HRP, a tyramide labeling was performed using the TSA kit (Molecular probes T20949) reagents while, if the secondary antibody was directly conjugated to Biotin, an incubation with streptavidin-HRP of 1h was performed and followed by Cy3 labeling (Perking Elmer TSA kit, NEL704A001KT). The sections were washed extensively before mounting in 80% glycerol/PBS. All the images (beside the one use for quantification, see below) were taken on the Leica SP8 confocal microscope with a 63x/1.30 Glycerol CORR objective and Hybrid detectors at 1024×1024 resolution in an 8-bit format. Brightness and contrast were adjusted using FiJi software and 3D reconstructions were performed with the Imaris software (Bitplane).

Antigen retrieval was performed using the TintoRetriever system from BioSB. The tissue was immerged in 10mM trisodium citrate buffer pH6.0 and brought to high temperature for 15 min using either the low pressure setting (106°C −110°C) or the high pressure setting (114°C −121°C) depending of the antibody (Table 1, supp. Methods). The samples were brought back gradually to RT before proceeding to the next step. For Frizzled3 staining we used a cycle of 20min at 95°C.

TUNEL assays were performed with an ApopTag Peroxidase *in situ* apoptosis detection kit (Millipore, S7100) and followed by immunofluorescence staining as described above.

### Image quantification

For quantification on the pancreatic epithelium, the entire pancreas was serially sectioned (8µm thick section) and the immunostained sections were imaged using a Leica DM5500 upright wide-field microscope with a 20X air objective. The positive pixels of the section stained for PDX1, SOX9 or β- +DAPI were measured on every 8th (E16.5), 6th (E14.5), 5th (E12.5) or every section (E10.5) using a customized Macro in Fiji software (Schindelin et al., 2012). The area obtained for the sampled sections were summed for each pancreas, which gives an estimation of the size of the epithelium. pHH3, CAS3 and TUNEL^+^ cells (matching a DAPI nucleus) were counted manually using Imaris software. For each pancreas sample, ratio between the number of counted cells (pHH3, CAS3 or TUNEL^+^) and the estimation of the size of the epithelium (based on SOX9, PDX1 or β-Catenin+DAPI areas of expression) were calculated and then normalized to the control littermates.

For the nervous system analysis, the whole embryo was serially sectioned (10µm thick section). After being stained for β-Catenin, DAPI and Caspase3 the sections were imaged using a Leica DM5500 upright wide-field microscope with a 20X air objective. Quantifications were made every 30^th^ section, which corresponds to 10 measurements per embryo. The neural tube was manually delimited based on β-Catenin staining using a free-hand-tool of Fiji. The area of neural tube was then calculated by measuring the positive pixels of β-Catenin+DAPI staining included in the drawn perimeter using the customized Macro of Fiji (Schindelin et al., 2012). Within the area of the neural tube we then calculated the area of Caspase3 expression with the same Macro. For each sample the ratio between the total area of Caspase3 expression/total neural tube area was calculated and normalized to control littermates.

### Western blot

E12.5 and E15.5 dorsal pancreata were snap-frozen and processed as described in^81^. Antibodies are listed in Table2 (supp. Methods). The blots were visualized using a Chemidoc MP (Bio-Rad) and Images were quantified based on pixel density using the ImageJ software package. The protein of interest was normalized to alpha tubulin level (internal control) then compared to the average value obtained for the control littermate.

### Pancreatic bud and sphere culture

For pancreatic bud culture, E10.5 dorsal bud epithelia were recovered, the mesenchyme was mechanically removed and the intact epithelium was seeded in 3D Matrigel culture. The buds were cultured in organoid medium (without Y27632)^25^ supplemented with either RhoA activator (CN03, Cytoskeleton Inc), Rock inhibitor (H-1152, #555550 Millipore) or an equal amount of DMSO for vehicle control. They were cultured for 5 to 7 days and the buds were imaged every day on a AF6000 Leica microscope with a 10x dry objective. In order to determine the working concentration of the drugs, a range of concentrations were initially tested (H1152: 0.25, 0.5, 1, 2.5, 5 and 10 µM; CNO3: 0.25, 0.5 and 1 µg/ml), the buds were collected after treatment and western blot for pMLC was performed to quantify the activity of RhoA/Rock pathway. 5 µM H1152 and 1 µg/ml CNO3 concentration were chosen as this respectively led to an average decrease of 68% and an average increase of 133% of pMLC protein amount while allowing an the epithelium to grow and branch. For the quantifications, the circumference of the growing epithelium was measured using Imaris software and exponential growth curves were generated for each bud using the GraphPad prism software. The doubling time of each individual growth curve was determined and used for statistical analysis. Normalization of the data (Fig. 4g) was done for each experimental condition by calculating the ratio (Δ) of the circumference (c) measured at each time point (t(x)) to the average circumference of the control littermate (C) at the same time point (Δ_t(x)_ = c_t(x)_ / C_t(x)_). Δ values were plotted against the time points of measurement and a linear regression was applied using GraphPad prism software. The slope of each individual line was determined and used for statistical analysis (Fig 4h). Of note, for the sake of clarity only the average growth curves or the average normalized linear regressions ares shown on the graphs for each experimental condition.

For the sphere cultures, E13.5 dorsal pancreata were isolated, the cells were dispersed, and seeded in Matrigel as described previously^25, 82^. Spheres were cultured for 4 days prior to live imaging and imaging was performed using either a SP8 Leica microscope with 20x glycerol objective for 5 hours (Litter1, 16 positions imaged for control and 22 positions imaged for *Ch-Vgl2* sample) or using a Zeiss LM780 with 20x dry objective for 3 hours (Litter2, 14 positions imaged for control, 13 for one *Ch-Vgl2* sample and 12 for the second *Ch-Vgl2* sample of the litter). Representative movies were made using Fiji software.

### Statistical analysis

Statistical analyses were performed with GraphPad 6 software packages and Microsoft Excel. Normalization to control littermates was obtain by making a ratio between the value obtained in the transgenics (or mutants) and the average value of controls present in the same litter, multiplied by 100 to convert to percentages. All statistical tests in the paper are unpaired non-parametric student t-tests. Normality of the data cannot be tested, we thus assume that the data have a Gaussian distribution. Results were indicated by the mean±SEM. Fold change calculations were made according to the formula: (final value-initial value)/initial value

