## Supplemental methods and figures for "Apical restriction of the planar cell polarity component VANGL in pancreatic ducts is required to maintain epithelial integrity"

**TABLE 1**: Technical details for sensitive immunohistochemistry

| Antibody | Antigen retrieval method | Blocking reagent | Secondary antibody | Amplification method |
|---|---|---|---|---|
| **DVL2** | none | TNB (a) | α-rabbit Biotinylated | streptavidin-HRP/Cy3 (a) |
| **FZD3** | 20min, 95˚C | BSA (b) | α-rabbit HRP | Tyr568 (b) |
| **PAR3** | Low pressure | BSA (b) | α-rabbit HRP | Tyr568 (b) |
| **PK2** | High pressure | BSA | α-rabbit cy3 FAB fragment | none |
| **pMLC** | none | BSA (b) | α-rabbit HRP | Tyr568 (b) |
| **Scribble** | Low pressure | BSA (b) | α-rabbit HRP | Tyr568 (b) |
| **VANGL** | High pressure | TNB (a) | α-rabbit Biotinylated | streptavidin-HRP/Cy3 (a) |

(a) Reagents from Perking Elmer TSA kit, NEL704A001KT (b) Reagents from Molecular probe TSA kit, T20949

**TABLE2**: list of antibodies used for immunohistochemistry

| Antigen | Species | Dilution | Reference |
|---|---|---|---|
| **Primary antibodies** | | | |
| β-Catenin | mouse | 1/1000 | BD transduction Lab 610153 |
| E-Cadherin | mouse | 1/100 | BD Transduction Lab C20820 |
| Cherry | rat | 1/1000 | Chromotek 5F8 |
| Cleaved-caspase 3 | rabbit | 1/200 | Cell Signaling Technology #9664 |
| Dishevelled2 | rabbit | 1/200 | Genetex GTX103878 |
| Frizzled3 | rabbit | 1/200 | Gift from J.Nathans |
| GFP | chicken | 1/1000 | Abcam ab13970 |
| Glucagon | guinea pig | 1/1000 | Linco 4031-01F |
| Insulin | guinea pig | 1/100 | Dako A0564 |
| Mucin1 | hamster | 1/1000 | Thermo Fisher Scientific HM-1630-P0 |
| Par3 | rabbit | 1/100 | Upstate Millipore 07330 |
| p-cJUN | rabbit | 1/1000* | Cell signaling technology #9164 |
| Pdx1 | goat | 1/1000 | Beta Cell Biology Consortium ab2027 |
| pHH3 | mouse | 1/100 | Cell signaling technology #9706 |
| α-PKC | rabbit | 1/500 | Santacruz sc-216 |
| pMLC | rabbit | 1/300, 1/1000* | Cell signaling technology #3674 |
| Prickle2 | rabbit | 1/1000 | Gift from T. Ohtsuka |
| Scribble | goat | 1/100 | Santacruz sc-c20 |
| Sox9 | rabbit | 1/500 | Chemicon AB5809 |
| Sox9* | rabbit | 1/2000 | Millipore AB5535 |
| Tubulin | rat | 1/10000* | Abcam ab6160 |
| Vangl1/2 | rabbit | 1/200 | Sigma Aldrich HPA025235 |
| Zo1 | mouse | 1/200 | Invitrogen 339100 |
| **Secondary antibodies** | | | |
| α-chick Al488 | goat | 1/1000 | Thermo Fisher A-11039 |
| α-goat Al488 | donkey | 1/1000 | Abcam ab150129 |
| α-goat Al488 | donkey | 1/1000 | Thermo Fisher A11055 |

| | | | |
|---|---|---|---|
| α-guinea pig Al488 | goat | 1/2500 | Trichem 345-FG-025/CF |
| α-guinea pig Al568 | goat | 1/800 | Thermo Fisher A11075 |
| α-hamster Al647 | goat | 1/1000 | Jackson Immuno Research 127-605-160 |
| α-hamster Biotinylated | goat | 1/500 | Jackson Immuno Research 127-065-160 |
| α-mouse Al488 | donkey | 1/1000 | Jackson Immuno Research 715-545-152 |
| α-mouse Al568 | donkey | 1/1000 | Thermo Fisher A10037 |
| α-mouse Al647 | donkey | 1/800 | Jackson Immuno Research 715-605-1500 |
| α-rabbit Biotinylated | donkey | 1/2000 | Thermo Fisher A16039 |
| α-rabbit HRP | donkey | 1/400 | Jackson Immuno Research 711-035-1520 |
| α-rabbit HRP* | donkey | 1/5000 | Dako P0048 |
| α-rabbit cy3 F(AB)$_2$ fragment | donkey | 1/500 | Jackson Immuno Research 711-166-152 |
| α-rabbit Al647 | donkey | 1/800 | Jackson Immuno Research 711-605-152 |
| α-rat cy3 | donkey | 1/1000 | Jackson Immuno Research 712-165-150 |
| α-rat Al647 | donkey | 1/500 | Jackson Immuno Research 712-605-153 |
| α-rat Al488* | donkey | 1/5000 | Jackson Immuno Research 712-545-153 |
| α-rat HRP* | donkey | 1/10000 | Jackson Immuno Research 712-035-153 |

*used for western blot

### Supplemental Figure 1

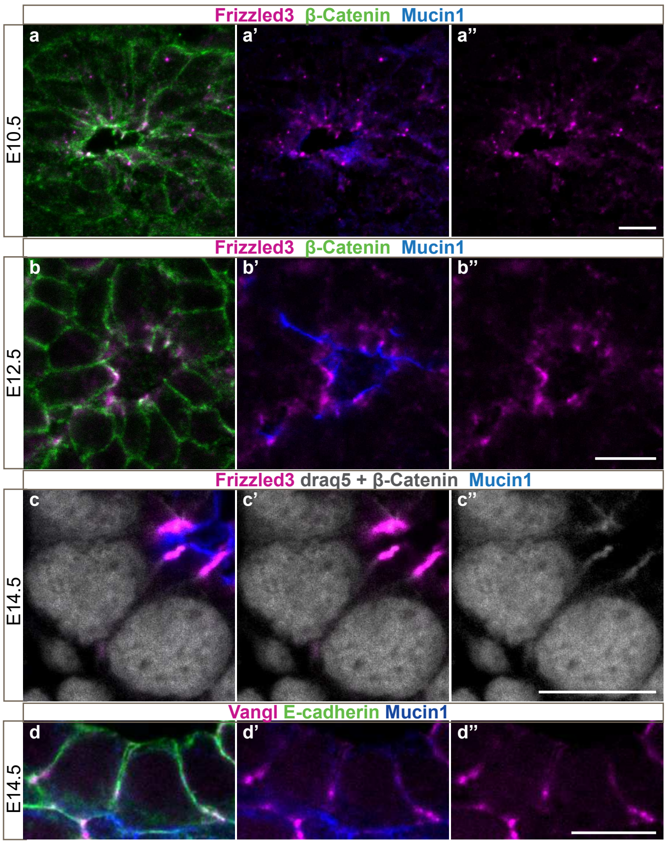


**FZD3 and VANGL expression domains become restricted to apical cell junctions during pancreas development**. Confocal images of embryonic pancreas sections stained for β-Catenin and Mucin1, labelling all and only apical membranes respectively. **a**, At E10.5, foci of FZD3 are detected around forming micro-lumen. **b**, By E12.5, FZD3 becomes enriched at the apical membrane but some cytoplasmic expression remains. **c-d**, At E14.5, both FZD3 and VANGL form stripes of staining on the lateral membranes, just below the apical membrane. Scale bar 10 µm

**Supplemental Figure 2**

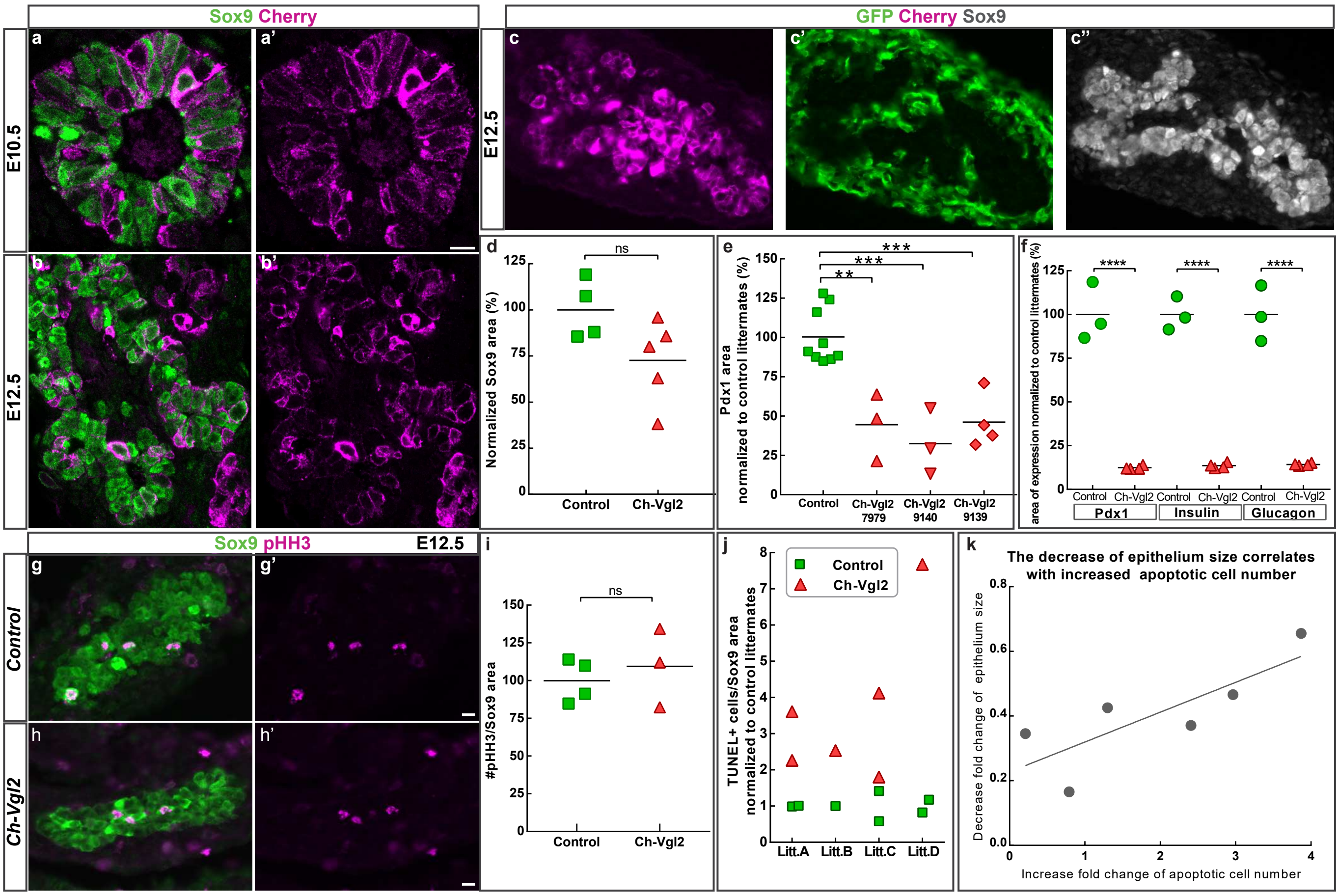


**Ectopic expression of VANGL on baso-lateral membranes leads to cell death and pancreatic hypoplasia**. **a-b**, Confocal images showing localization of Cherry-VANGL2 fusion protein in double-transgenic *Pdx1-Cre; Cherry-Vangl2* (*Ch-Vgl2*) on E10.5 and E12.5 pancreatic sections. The fusion protein initially located at the cell cortex of most of the epithelial cells (SOX9$^{+}$) (a) becomes restricted to the cell membranes (b). **c**, Recombination of the transgene in the pancreatic progenitors (*Pdx1-Cre* driver) allows the detection of Cherry-VANGL2 fusion protein (magenta) in the epithelium (SOX9$^{+}$, grey) while the surrounding mesenchymal cells, where recombination does not occur, remain GFP$^{+}$ (green). **d**, SOX9 area quantified on E10.5 embryonic sections of control (green) and *Ch-Vlg2* (pink) pancreas shows no significant change of the epithelium size in *Ch-Vgl2*. p-value=0.08 by t-test. **e**, PDX1 area quantified on E12.5 sections of control (green) and double transgenic (pink) pancreata isolated from 3 different *Ch-Vgl2* mouse lines shows a significant decrease of the epithelium size in all *Ch-Vgl2* lines. (**) p-value= 0.001 (***) p-value= 0.0003 by t-test. **f**, PDX1, Insulin and Glucagon expression areas were quantified on E16.5 pancreas sections. The pancreatic progenitors (PDX1$^{+}$) as well as the endocrine cells (Insulin$^{+}$ and Glugagon$^{+}$) expression areas are significantly decreased in the *Ch-Vgl2* (pink) compared to the control siblings (green), revealing a hypoplasia of the *Ch-Vgl2* pancreas. (****) p-value ≤ 0.0001 by t-test. **g-i**, pHH3 immunostaining on E12.5 pancreas sections shows no change in proliferating cell numbers (pHH3$^{+}$, magenta) within the pancreatic progenitors domain (SOX9$^{+}$, green). **i**, quantification of pHH3$^{+}$ cells number/ epithelium area (SOX9$^{+}$) ratio based on the staining presented in g-h. p-value= 0.6 by t-test. **j**, Graph showing the quantification of the TUNEL assay presented in figure 2h. The apoptotic cell (TUNEL$^{+}$) numbers include dead cells located within the pancreatic epithelium (SOX9$^{+}$) and those lying just in the periphery of the epithelium among the mesenchymal cells (arrow in Fig. 2h). The ratios, *i.e* number of TUNEL$^{+}$ cells/ epithelium area, are normalized against the control littermates and data are presented by litters. Including the TUNEL$^{+}$ cells located in the periphery of the epithelium in the quantification performed in Fig. 2h exacerbates the phenotype observed in the *Ch-Vlg2* pancreas, showing a 3 fold increase in apoptosis in the *Ch-Vgl2* compared to control littermates. p-value=0.007 by t-test. **k**, Correlation graph showing for each individual *Ch-Vgl2* pancreas (one dot) collected at E12.5 the fold change in the size of the epithelium (absolute value obtained from quantifications presented in Fig. 2e are plotted in y) and the fold change in the number of apoptotic cells (x value, obtained from quantification presented in Fig. 2h) compared to the control littermate. A Pearson correlation test applied on this dataset reveals that the extent of hypoplasia positively correlates with the increase of apoptosis (Pearson r=0.8, p-value ≤0.05). (**d-f, i-k**) For all the graphs, each dot represents one pancreas and the data are normalized against control littermates. Scale bar= 10 µm.

### Supplemental Figure 3

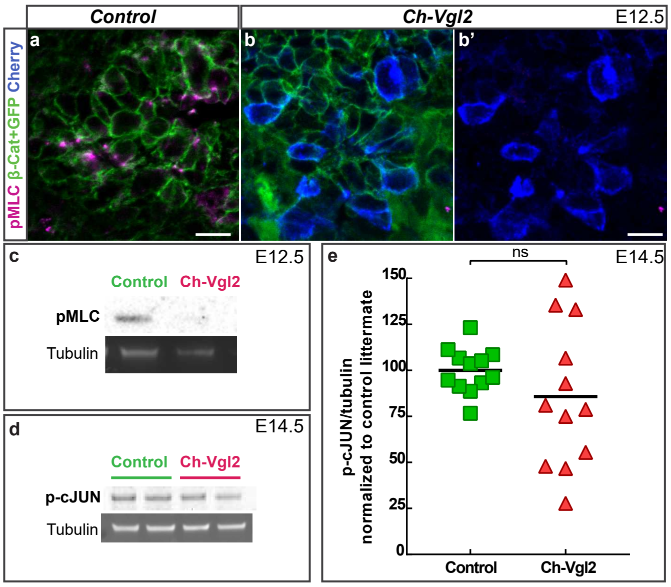


**VANGL misexpression affects ROCK but not JNK pathway activity**. **a-b**, confocal images showing expression of pMLC on E12.5 pancreas sections. Note the disappearance of pMLC apical foci in the *Ch-Vlg2* section (b). **c**, representative image of the pMLC western blot performed on E12.5 pancreas. The pMLC band becomes almost undetectable in the *Ch-Vgl2*. **d**, representative image of the p-cJUN western blot performed on E14.5 pancreas and used for the quantification in e. **e**, p-cJUN/Tubulin ratios are normalized against the control littermates, each dot corresponds to one pancreas. There is no significant change in the p-cJUN protein levels in the transgenic population compared to the control. T-test, p-value = 0.3. Scale bar 10 µm.

### Supplemental Figure 4

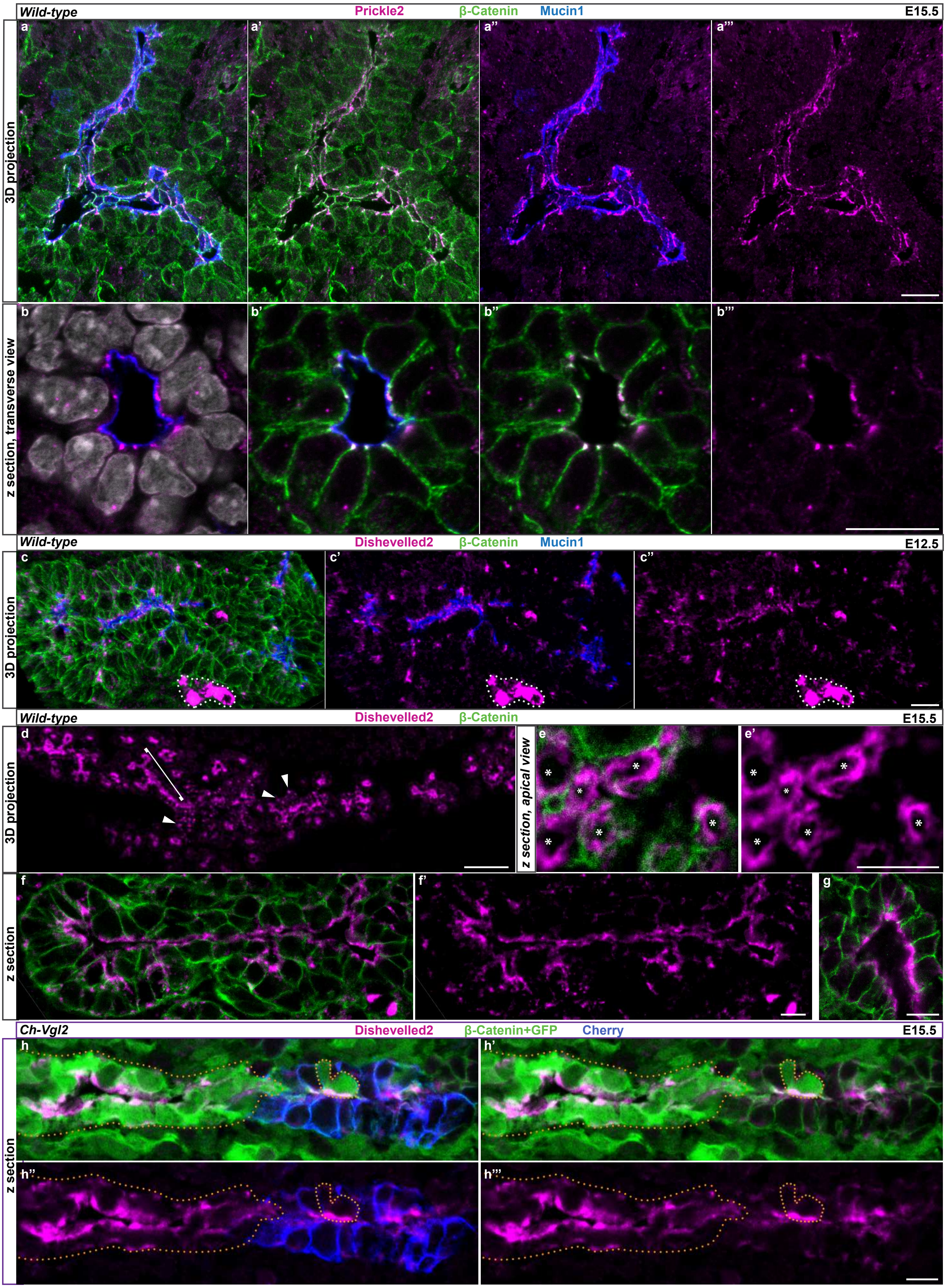


**PK and DVL cellular localization in the developing pancreas**. Confocal images of embryonic pancreas sections stained for β-Catenin labeling all the membranes, Mucin1 labeling apical membranes, Prickle2 (PK2) (a-b) and Dishevelled2 (DVL2) (c-h). **a-b,** Representative images of PK2 expression in a general view of the epithelium (a) or a close-up of a transverse section of a duct (b) showing that expression of PK2 is restricted to the apical cell junctions in the E14.5 pancreas. **c**, Representative images of DVL2 expression in an overview of the epithelium at E12.5. DVL2 protein is enriched at the apical side of the duct highlighted by Mucin1 labelling (blue). A strong cytoplasmic expression of DVL2 is also detected in the delaminating endocrine cells (dashed line) **d**, low magnification (20X) of a section through an E15.5 dorsal pancreas. The continuous DVL2 staining (bracket for example) underlines the apical side of ducts while the more dotted labeling (arrow for example) marks the endocrine cells. **e**, Apical view of ductal cells (one* per cell) in a longitudinal section of a large duct showing rings of DVL2 staining at the apical cortex. **f-g**, close-up of d showing the apical localization of DVL2 in a duct. Note that DVL2 staining is continuous at the apical membrane (g) conversely to the other core components that are restricted to apical cell junctions (VANGL: Fig. 1b, FZD3: Fig. 1d and PK2: b). **h**, Longitudinal section of a E15.5 *Ch-Vgl2* duct showing that DVL2 expression (magenta) is decreased in the Cherry+ cells (blue) compared to the adjacent cells that are not expressing the Cherry-VANGL2 fusion protein (green and highlighted by orange dots and arrowheads). Scale bar = 10 µm, except d= 100 µm.

### Supplemental Figure 5

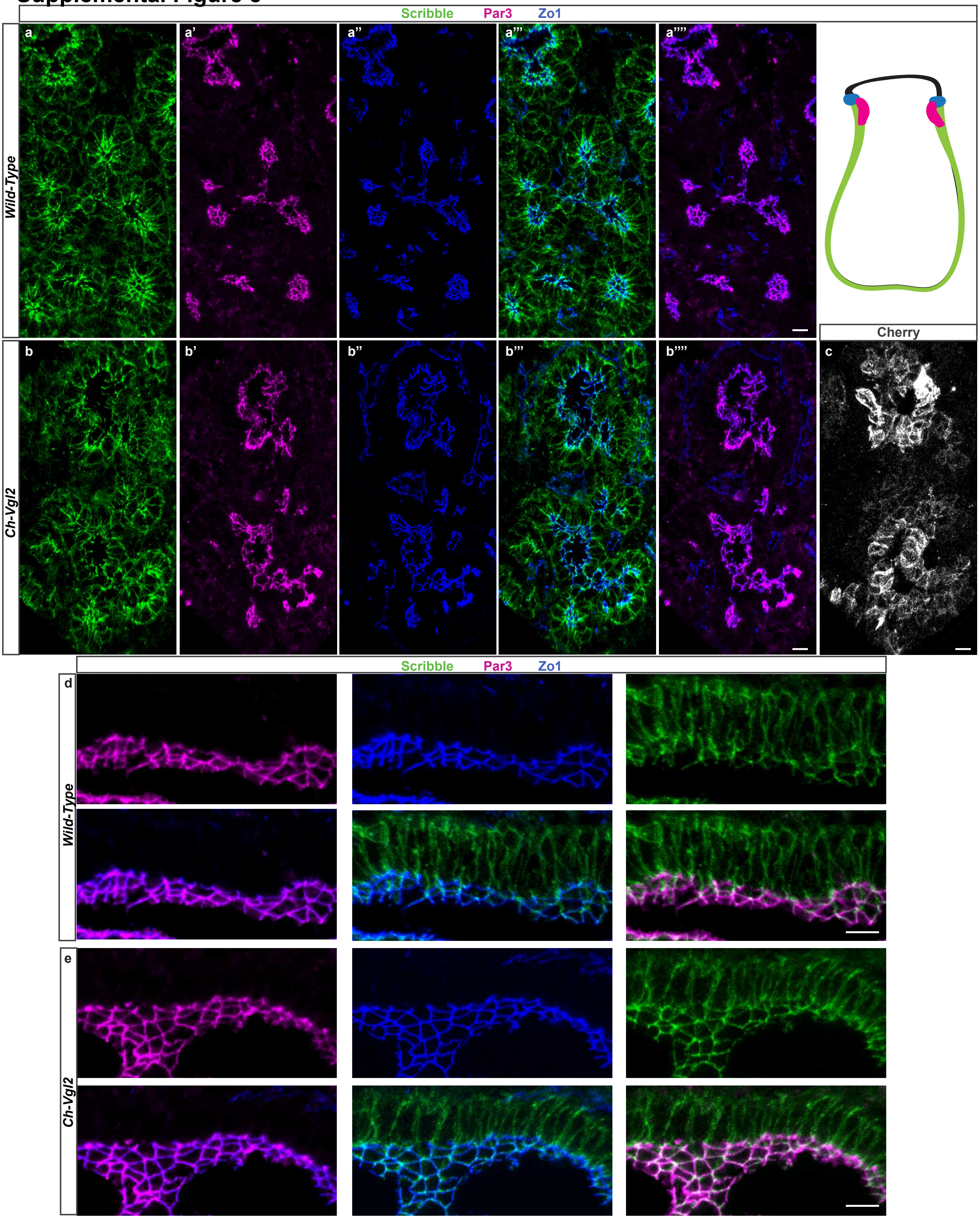


**The expression of apico-basal components is not affected following ectopic VANGL protein expression.** Confocal images of E15.5 pancreatic sections, showing the expression pattern of Scribble (green), PAR3 (magenta) and ZO1 (blue) in a Cherry-VANGL2$^{+}$ area (b,e). All images are 3D-projections **a-c**, General view of the pancreatic epithelium. In wild-type pancreas (**a**) Scribble is expressed at the baso-lateral membrane of epithelial cells and is enriched around the adherent junctions. The tight junction maker ZO1 is expressed slightly more apically than Scribble while the PAR3 expression domains overlaps with both ZO1 and the most apical expression of Scribble (scheme). This expression pattern is conserved in the double transgenics (**b**) in areas where Cherry-VANGL2 fusion protein is strongly expressed (**c**). **d-e**, zoom in a section of a large pancreatic duct allowing an apical view of ductal cells. Scribble, PAR3 and ZO1 relative expression domains are unchanged in *Ch-Vgl2* compared to controls. Scale bar=10 µm
