## Supplementary material for "Apical restriction of the planar cell polarity component VANGL in pancreatic ducts is required to maintain epithelial integrity": Movie legends

**Movie1**

**VANGL expression is planar polarized in the pancreatic duct**. Whole mount staining for VANGL (red), β-Catenin (green) and Mucin1 (grey) showing a large duct. At time 27-30s note the chevron like localization of VANGL at the apical surface of the cells, on membranes perpendicular to the longitudinal axis of tubes.

**Movie2-7**

**Epithelial exit and destruction of epithelial integrity upon ectopic VANGL expression.** Pancreatospheres generated from E13.5 *Ch-Vgl2* (M2, M4, M5 and M7) and control littermates (M3 and M6) epithelium were imaged every _῀_12 min for 5 hours (M2-M4, M7) or 3 hours (M5-M6) in brightfield (grey), to visualize the whole epithelial layer, and with fluorescent filter to visualize GFP- (green) and Cherry- (magenta) positive cells. M2-M6 are Optical z sections through a live sphere while M7 is a 3D projection. **M2-M4**, Epithelium thickening and cell delamination resulting in loss of epithelium integrity was observed in the *Ch-Vgl2* spheres (M2) but not in the control littermates (M3). Eventually, the *Ch-Vgl2* spheres collapsed entirely (M4). **M5-M6**, similar phenotypes were observed in shorter movies (3 h). While the size of the lumen remained roughly constant in the control spheres (M6), the lumen of *Ch-Vgl2* spheres decreased and ultimately disappeared (M5). **M7**, *Ch-Vgl2* epithelium loss of integrity is also visible in 3D projection, showing the epithelial cell wobbling and rounding.
